# Stimulation of locus coeruleus inputs to the frontal cortex in mice induces cell type-specific expression of the *Apoe* gene

**DOI:** 10.1101/2024.07.22.604695

**Authors:** Genevieve E. Craig, Lizbeth Ramos, Samuel R. Essig, Nicholas J. Eagles, Andrew E. Jaffe, Keri Martinowich, Henry L. Hallock

## Abstract

Deficits in attention are common across a range of neuropsychiatric disorders. A multitude of brain regions, including the frontal cortex (FC) and locus coeruleus (LC), have been implicated in attention. Regulators of these brain regions at the molecular level are not well understood, but might elucidate underlying mechanisms of disorders with attentional deficits. To probe this, we used chemogenetic stimulation of neurons in the LC with axonal projections to the FC, and subsequent bulk RNA-sequencing from the mouse FC. We found that stimulation of this circuit caused an increase in transcription of the *Apoe* gene. To investigate cell type-specific expression of *Apoe* in the FC, we used a dual-virus approach to express either the excitatory DREADD receptor hM3Dq in LC neurons with projections to the FC, or a control virus, and found that increases in *Apoe* expression in the FC following depolarization of LC inputs is enriched in GABAergic neurons in a sex-dependent manner. The results of these experiments yield insights into how *Apoe* expression affects function in cortical microcircuits that are important for attention-guided behavior, and point to interneuron-specific expression of *Apoe* as a potential target for the amelioration of attention symptoms in disorders such as attention-deficit hyperactivity disorder (ADHD), schizophrenia, and Alzheimer’s disease (AD).

**Significance Statement:** Identifying patterns of gene expression in specific brain circuits is an important first step toward developing treatments for cognitive and behavioral symptoms that rely on those circuits. In this paper, we describe a transcriptome-scale motif in one such circuit - neurons in the LC that project to the FC. This circuit has been implicated in attention, and attentional deficits are common across many neuropsychiatric disorders, suggesting that targeting this circuit could have therapeutic potential for ameliorating attentional symptoms in these disorders. We further explored one of the top differentially expressed genes, *Apoe,* to identify how it is expressed in distinct cell types following stimulation of this circuit, paving the way for spatially- and genetically-specific targeting of this gene in attention.

## Introduction

Deficits in attention are common across a range of neuropsychiatric disorders, including attention deficit hyperactivity disorder (ADHD), major depressive disorder (MDD), and schizophrenia (Braff, 1993; Hooks et al., 1994; Paelecke-Habermann et al., 2005). The anterior cingulate cortex (ACC), located on the medial surface of the frontal lobes in humans, is heavily involved in attention - for example, the ACC is highly active during sustained attention, defined as a state of readiness for detecting unpredictable events (MacDonald et al., 2000). The ACC is also involved in conflict monitoring, or the allocation of attention based on conflicting signals (Carter et al., 1998). For example, Fallgatter et al. (2002) showed that the ACC exhibits greater activity during response inhibition while humans perform the continuous performance task (CPT) of attention, further supporting the role of the ACC in conflict monitoring. The ACC also modulates cognitive control in other areas of the brain, such as the prefrontal cortex (PFC) (Kerns et al., 2004), potentially via neuromodulatory systems like the locus coeruleus (LC). The LC supplies cortical norepinephrine (NE) and is reciprocally connected to the ACC in primates (Joshi & Gold, 2022) and rodents (Jodoj et al., 1998; Pudovkina et al., 2001). LC-NE activation influences ACC activity patterns and affects attention (Dahl et al., 2020), implicating the LC as a driver of cortical function in attention-guided behavior. Deficits in ACC and LC function are also present in disorders in which attention deficits are a common cognitive symptom, such as schizophrenia, ADHD, and MDD (Kerns et al., 2005; del Cerro et al., 2020; Shirama et al., 2020).

Translating research on the neurobiology of attention to rodents has been difficult, in part because identifying regions that are homologous to human cortical areas is not straightforward. Recent research has provided strong evidence that the area commonly referred to as the medial prefrontal cortex (mPFC) in rodents is actually more anatomically homologous with the human cingulate cortex (Laubach et al., 2018; van Heukelum et al., 2020), providing an avenue for comparison of attention-related activity between species. In support of this, many studies in rodents have shown that activity in this area of the brain (referred to here as the frontal cortex, or FC) is strongly related to attention-specific behavior (Birrell & Brown, 2000; Passetti et al., 2002; Fisher et al., 2020). Furthermore, catecholamine signaling in the rodent FC is involved in attention (Berridge & Spencer, 2016), and drugs that modulate catecholaminergic function often affect attention in rodents (Mar et al., 2017; Caballero-Puntiverio et al., 2019), suggesting that brain areas like the LC drive function in cortical areas to promote rodent attention as well. The FC and LC functionally interact during a rodent version of the CPT (Hallock et al., 2024), further corroborating the idea that the LC and FC work together during attention-guided behavior across mammalian species.

Although specific brain circuits, like the one between the LC and FC, have been implicated in attention, little research has been done to understand the molecular drivers of function in these circuits. Gene expression in cortical tissue is heterogeneous between brain regions and cell types in both humans (Darmanis et al., 2015; Maynard et al., 2021) and rodents (Zeisel et al., 2015), and these patterns of gene expression change in response to behavioral experience (Cho et al., 2016; Mukherjee et al., 2018), suggesting that unique cell type-specific patterns of gene expression could be used as markers of circuit function. These results also raise the possibility that cells embedded in specific circuits could be non-invasively accessed by manipulating genes selectively expressed in those cells, paving the way for gene-targeted treatments for cognitive symptoms in neuropsychiatric disorders, such as deficits in attention. In order to uncover gene expression motifs in one such circuit, we used chemogenetic targeting to activate neurons in the LC that send direct axonal projections to the FC in mice. We subsequently used bulk RNA-sequencing and single-molecule *in situ* hybridization to look at cell type-specific transcription in the FC in response to stimulation of LC inputs. We found that, among other genes, the gene encoding for the apolipoprotein-E protein (*Apoe*) was enriched in FC tissue following circuit activation. This increase in *Apoe* transcription was driven by expression in GABAergic neurons, and this effect was more robust in female mice.

## Materials and Methods

### Subjects

For the RNA-sequencing experiment, we used a cohort of 8 male (4 experimental and 4 control) wild-type c57bl/6j mice (Jackson strain 000664). For the RNAscope experiments, we used a cohort of 12 female (6 experimental and 6 control) and 12 male (6 experimental and 6 control) wild-type c57bl/6j mice (24 total mice). Mice were group-housed (3-5 animals per cage) with *ad libitum* access to both food and water. The colony room was temperature and humidity controlled on a 12 h light/dark cycle. All experiments were performed during the light cycle. At time of surgery, animals were roughly 90 to 120 days of age. All procedures were in accordance with the Institutional Animal Care and Use Committees of [Author University] (mice used in the RNA-sequencing experiment) and [Author University] (mice used in RNAscope experiments).

### Surgical and Extraction Procedures

For all experiments, mice were anesthetized with isoflurane (1-2.5% oxygen) and then placed into a stereotaxic frame (Kopf Instruments, Tujunga, CA). An incision was made along the midline of the scalp, the skull was leveled, and bregma was identified. Holes were drilled in the skull above the target brain regions (LC: −5.4 mm AP from bregma, ±0.9 mm ML from the midline; FC: +1.7 mm AP from bregma, ±0.3 mm ML from the midline), and an automated infusion pump (World Precision Instruments, Sarasota, FL) was used to inject the viruses at 3 nl/sec for a total volume of 600 nl/hemisphere in the FC (1.7 mm ventral to the surface of the brain) and 300 nl/hemisphere in the LC (3.0 mm ventral to the surface of the brain). Measurements for these brain regions were obtained from the Paxinos and Franklin mouse brain atlas (Paxinos & Franklin, 2019). For the RNA-sequencing experiment, a retrograde virus encoding Cre-recombinase (AAVrg-hSyn-Cre; Addgene catalog # 105553-AAVrg) was injected into the FC, and a virus coding for the Cre-dependent expression of an excitatory DREADD receptor (AAV8-hSyn-DIO-hM3Dq-mCherry; Addgene catalog # 44361-AAV8) was injected into the LC (Fig. 1A). For RNAscope experiments, AAVrg-hSyn-Cre was injected into the FC, and either a virus coding for the Cre-dependent expression of an excitatory DREADD receptor (AAV8-hSyn-DIO-hM3Dq-mCherry; experimental group) or a control virus (AAV8-hSyn-DIO-mCherry; control group; Addgene catalog # 114472-AAV8) was injected into the LC (Fig. 2A). For the RNA-sequencing experiment, we waited 4-5 weeks for the virus to infect a sufficient number of cells, and then gave intraperitoneal (IP) injections of either clozapine-N-oxide (CNO; 2.5 mg/kg; experimental group; Tocris Biosciences catalog # 4936), or vehicle (filtered 1x phosphate-buffered saline (PBS); control group) into the mice. Vehicle injections were matched to the volume of CNO injections in experimental mice. Because CNO exerts its peak effects after ∼30 minutes, and peak expression of many immediate early genes (IEGs) is typically seen around 90 minutes after stimulation (Barry & Commins, 2017), we killed the mice 120 minutes following injections. Following cervical dislocation, the brains of the mice were extracted, the medial wall of the FC from both hemispheres was dissected with a brain block and razor blades on wet ice, and each hemisphere was flash-frozen in 2-methylbutane and stored in a 1.5 ml Eppendorf centrifuge tube at −80 degrees C. For RNAscope experiments, we again waited 4-5 weeks for the virus to infect a sufficient number of cells, gave all mice CNO injections (2.5 mg/kg; IP), and killed the mice 120 minutes following injections. We then extracted the brains, flash-froze them in 2-methylbutane, and stored them at −80 degrees C.

**Fig. 1:**
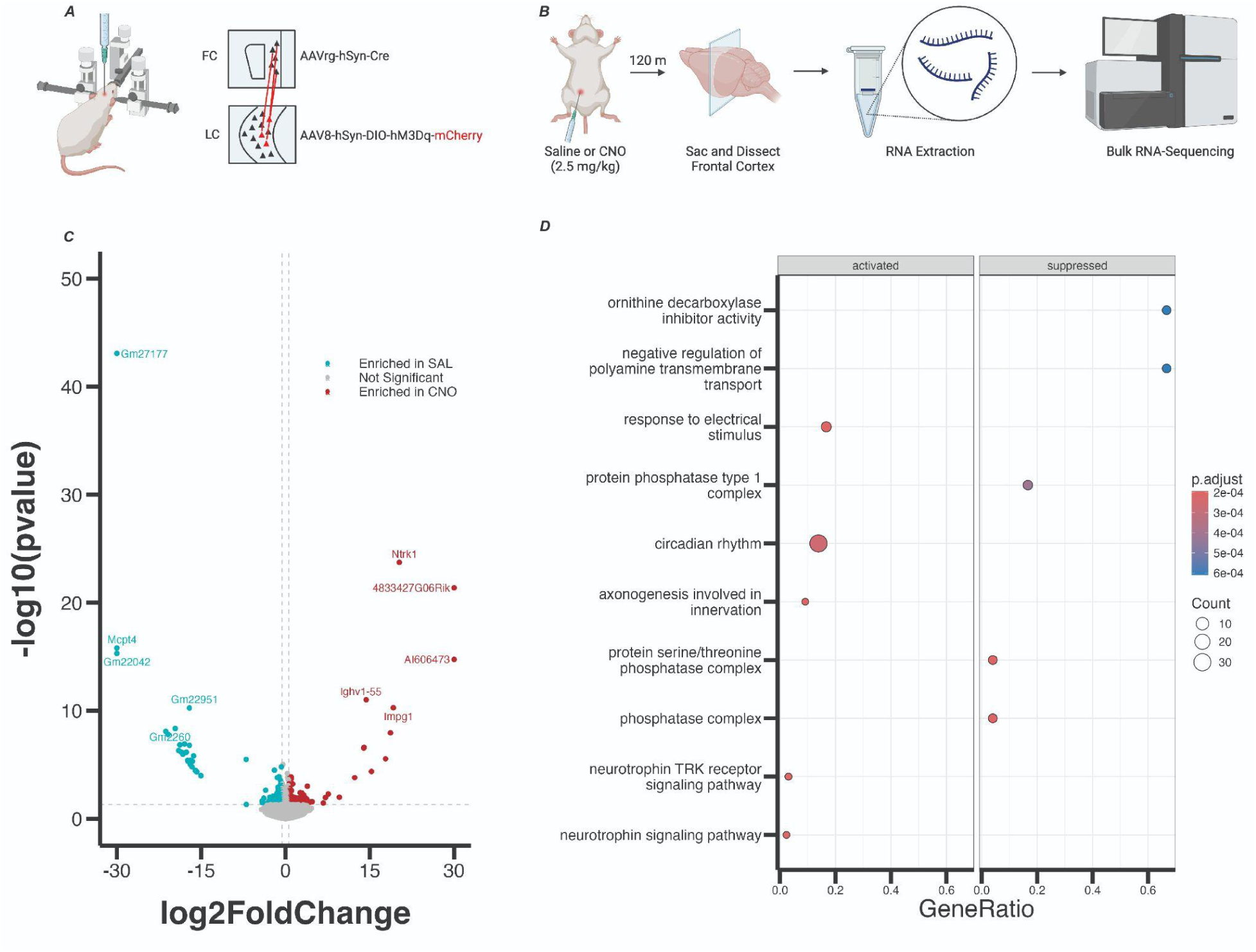
(A) Schematic showing viral injection strategy for chemogenetic activation of frontal cortex (FC)-projecting neurons in the locus coeruleus (LC). (B) Experimental design for identification of enriched gene sets in the FC following chemogenetic activation of FC-projecting LC neurons. (C) Volcano plot showing genes that were enriched in FC tissue in the experimental (CNO-receiving) group of mice (red dots), and genes that were enriched in FC tissue in the control (saline-receiving) group of mice (blue dots) based on differential expression analysis. The top 10 differentially-expressed genes (based on FDR-adjusted p-value) are labeled. (D) Dot plot showing gene ontology terms that contained differentially-expressed gene sets for experimental mice (left column) and control mice (right column). The top ten gene ontology terms by FDR-adjusted p-value are listed on the y-axis.

**Fig. 2:**
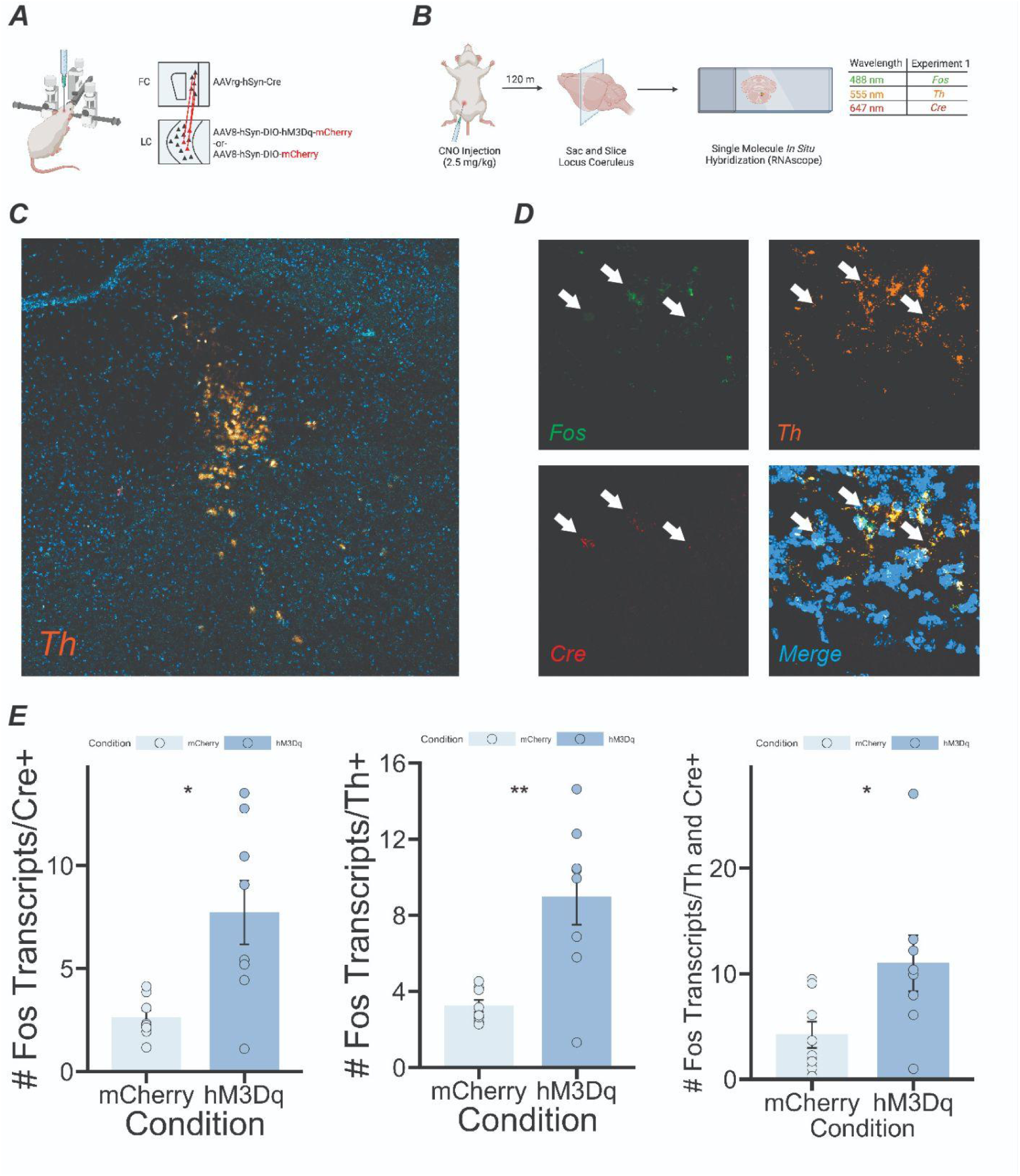
(A) Schematic showing viral injection strategy for activation of FC-projecting neurons in the LC for RNAscope experiments. (B) Experimental design (C) Confocal image showing tyrosine hydroxylase (Th) expression in the locus coeruleus (LC). (D) Confocal image at 40x magnification showing expression of Fos, Th, and Cre in LC tissue. (E) In mice that had the DREADD receptor, Fos expression was increased in Cre+ cells (left panel - t(7.786) = 3.2078, p = 0.01292), Th+ cells (middle panel - t(7.6345) = 3.8082, p = 0.005647), and cells that were both Th and Cre+ (right panel - t(9.9693) = 2.3047, p = 0.04398) compared to mCherry controls. Error bars = SEM.

### Bulk RNA-Sequencing

Total RNA was isolated and extracted from the tissue using TRIzol (Life Technologies, Carlsbad, CA), purified using RNeasy minicolumns (Qiagen, Valencia, CA; catalog # 74104), and quantified using a NanoDrop spectrophotometer (Agilent Technologies, Savage, MD). The Nextera XT DNA Library Preparation Kit was used to generate sequencing libraries according to manufacturer instructions, and samples were sequenced on a HiSeq2000 (Illumina). Reads were aligned to the mm10 genome using the HISAT2 splice-aware aligner (Kim et al., 2015), and alignments overlapping genes were counted using featureCounts version 1.5.0-p3 relative to Gencode version M11. Differential expression analyses were performed on gene counts using the voom approach (Law et al., 2014) in the limma R/Bioconductor package (Ritchie et al., 2015) using weighted trimmed means normalization factors with condition (CNO vs. saline) as the main outcome of interest, and adjusting for the exonic mapping rate. Multiple testing correction was performed using the Benjamini-Hochberg approach to control for the false discovery rate (FDR). Gene ontology (GO) analyses were performed using ENSEMBL gene IDs with the clusterProfiler R Bioconductor package (Ashburner et al., 2000; Yu et al., 2012).

### Single-Molecule Fluorescent In Situ Hybridization

To perform single-molecule *in situ* hybridization (RNAscope) on LC and FC tissue, we took coronal sections of the FC and LC (16 μm) on a cryostat (Leica, Nussloch, Germany), mounted them onto slides, and performed the RNAscope protocol using the fluorescent multiplex V2 kit from ACDBio (catalog # 323110). Specifically, tissue sections were briefly fixed with a 10% neutral buffered formalin solution at room temperature, and subjected to serious dehydration with ethanol. We then pretreated the sections with protease IV and hydrogen peroxide, and incubated the slides at 40 degrees C with a combination of three probes. For FC tissue, these probes were as follows - channel 1: *Gfap*, channel 2: *Rbfox3*, channel 3: *Apoe* for experiment 1; channel 1: *Slc17a7*, channel 2: *Gad1*, channel 3: *Apoe* for experiment 2; channel 1: *Sst*, channel 2: *Pvalb*, channel 3: *Apoe* for experiment 3. For LC tissue, these probes were as follows - channel 1: *Fos*, channel 2: *Th*, channel 3: *Cre*. After incubation, we applied amplification buffers for each channel, and opal dyes (520 nm for channel 1; 570 nm for channel 2; 690 nm for channel 3; Akoya Biosciences, catalog #’s FP1487001KT, FP1488001KT, and FP1497001KT) in order to fluorescently label each transcript. Lastly, we stained the sections with DAPI to demarcate the nuclei of the cells. We then took z-stacked images of the FC (four sections per slide, one image per section, four images per mouse total) and LC slides (one section per slide, one image per section, one image per mouse total) using a Zeiss LSM800 confocal microscope. For analysis, we used a MATLAB program to quantify transcript expression in each image (*dotdotdot*; Maynard et al., 2020). Specifically, we used the ‘CellSegm’ toolbox to perform nuclear segmentation in x, y, and z-dimensions to define regions of interest (ROIs) based on DAPI expression, and watershed analysis to identify distinct transcripts (individual dots) in each of the three microscope channels corresponding to an opal dye. We then co-localized each identified transcript with an identified nucleus (ROI). Transcripts that were not classified by the program as being co-localized with a nucleus were not used for analysis. Background noise, which could result from bleed-through from adjacent wavelengths in the gene channels, was eliminated using the ‘imhmin’ function. This function suppresses all of the minima in the grayscale image whose depth is less than the standard deviation of the image. We used a cutoff of 5 transcripts to categorize a cell as either *Rbfox3* or *Gfap*-expressing, *Gad1* or *Slc17a7*-expressing, and *Pvalb* or *Sst*-expressing, respectively

### Statistical Analysis

For RNAscope results, we used multiple two-way analyses of variance (ANOVAs) to compare gene expression between sex and experimental condition (DREADD vs. mCherry controls). We also used Welch’s two-sample t-tests to compare gene expression in the LC between DREADD and mCherry controls. An alpha level of 0.05 was used to determine statistical significance. All tests were performed with the R programming language (‘aov’ and ‘t.test’ functions).

## Results

### RNA-Sequencing

We identified 96 genes that were differentially expressed in FC tissue between our CNO and saline groups (genes that had an FDR-adjusted p-value < 0.05). Of these genes, 82 were enriched in the saline group, and 14 were enriched in the CNO group (Fig. 1c). Gene ontology (GO) analysis revealed that genes enriched in the CNO group are involved in circadian rhythms, axonogenesis, and neurotrophin signaling, suggesting that stimulation of LC inputs to the FC induces signaling pathways involved in plasticity (Fig. 1d). Of the genes that were enriched in the CNO group, the *Apoe* gene (log2FoldChange = 0.34, FDR-adjusted p-value = 0.0342) stood out due to its involvement in attention in humans (Parasuraman et al., 2002; Rusted et al., 2013). A complete list of differentially-expressed genes is available in the extended data (Table 1-1).

### Single-Molecule In Situ Hybridization

In order to verify our RNA-sequencing results, and determine cell type-specific expression of *Apoe* in FC tissue following depolarization of LC inputs, we replicated our RNA-sequencing experiment with several important changes. First, we used both male and female mice in order to determine whether sex differences in *Apoe* expression levels might be present, as previous research has demonstrated sex differences in mouse locus coeruleus anatomy and function (Bangasser et al., 2011; Bangasser et al., 2016). Second, we employed an experimental design in which all mice received CNO injections, as the presence of CNO alone could have accounted for the increase in *Apoe* transcription that we observed in our RNA-sequencing data.

Firstly, to verify that our viral targeting strategy increased neuronal activity in LC neurons, we took coronal sections of the LC and quantified *Fos* expression in neurons expressing tyrosine hydroxylase (*Th*-expressing neurons) and neurons expressing Cre-recombinase (*Cre*-expressing neurons) (Fig. 2c and Fig. 2d). Due to the difficulties inherent in slicing LC tissue, we were only able to obtain usable slices from 16 total mice (12 male, 4 female; 8 experimental, 8 control). We found evidence that *Th* and *Cre*-expressing neurons in the LC were robustly activated following CNO injections in the experimental group, as *Fos* transcription was increased in *Cre*-expressing (*t*(7.786) = 3.2078, *p* = 0.01292), *Th*-expressing (*t*(7.6345) = 3.8082, *p* = 0.005647), and both *Cre* and *Th*-expressing (*t*(9.9693) = 2.3047, *p* = 0.04398) neurons in the LC of experimental mice compared to mice in the control group (Welch’s t-tests; Fig. 2e).

We next looked for evidence of *Apoe* expression in astrocytes and neurons in the FC following chemogenetic stimulation of LC inputs (Fig. 3b), and found that neither number of *Apoe* transcripts nor percentage of cells expressing *Apoe* significantly differed between group or sex in astrocytes (*Gfap*-expressing cells; Fig. 3c), but that both number of transcripts and percentage of cells expressing *Apoe* were significantly increased in putative neurons (*Rbfox3*-expressing cells; *F*(1,84) = 10.498, *p* = 0.00171, main effect of group for number of *Apoe* transcripts; *F*(1,84) = 40.078, *p* = 1.14e-8, main effect of condition for percentage of neurons expressing *Apoe*). Additionally, the number of *Apoe* transcripts expressed in *Rbfox3*-expressing cells was significantly higher in females, compared to males (*F*(1,84) = 15.777, *p* = 0.00015, main effect of sex; Fig. 3d).

**Fig. 3:**
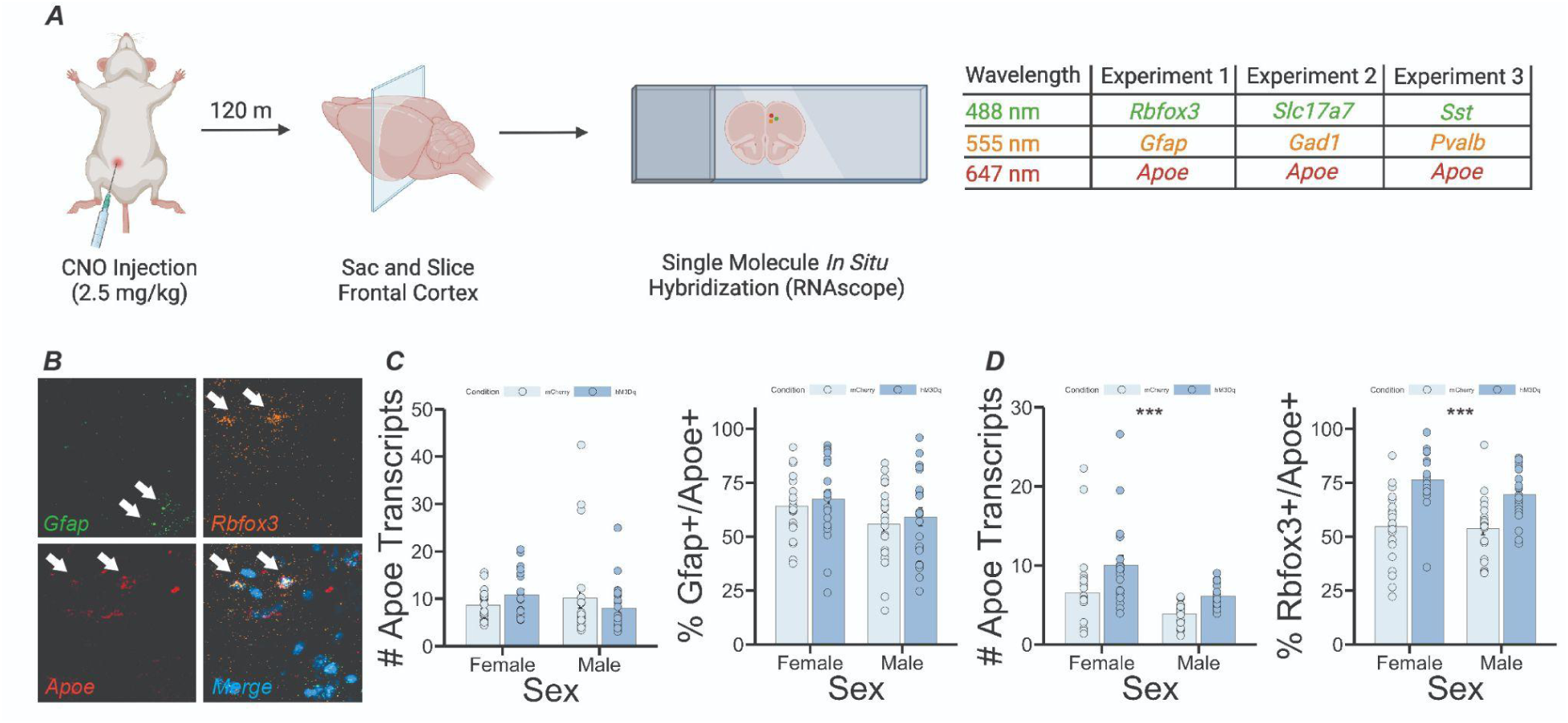
(A) Experimental design (B) Confocal image showing expression of Gfap, Rbfox3, and Apoe in FC tissue. (C) No significant differences in Apoe expression in astrocytes were observed between conditions or sexes. (D) Apoe expression significantly increased in neurons following LC depolarization (F(1,84) = 15.777, p = 0.00015, main effect of sex, F(1,84) = 10.498, p = 0.00171, main effect of condition for # transcripts, and F(1,84) = 40.078, p = 1.14e-8, main effect of condition for % neurons).

To determine whether neuronal increases in *Apoe* were predominantly in excitatory (glutamatergic) or inhibitory (GABAergic) neurons, we next looked at *Apoe* expression in *Slc17a7*-expressing cells (*Slc17a7* encodes the glutamate transporter protein), and *Gad1*-expressing cells in FC tissue (Fig. 4a). We found that neither number of *Apoe* transcripts, nor percentage of cells expressing *Apoe* significantly differed between group or sex in putative excitatory neurons (Fig. 4b), but were both significantly increased in putative inhibitory neurons in the experimental group compared to the control group, irrespective of biological sex (*F*(1,84) = 9.154, *p* = 0.00329, main effect of group for number of *Apoe* transcripts; *F*(1,84) = 12.898, *p* = 0.000553, main effect of group for percentage of *Gad1*-expressing neurons co-expressing *Apoe*; Fig. 4c).

**Fig. 4:**
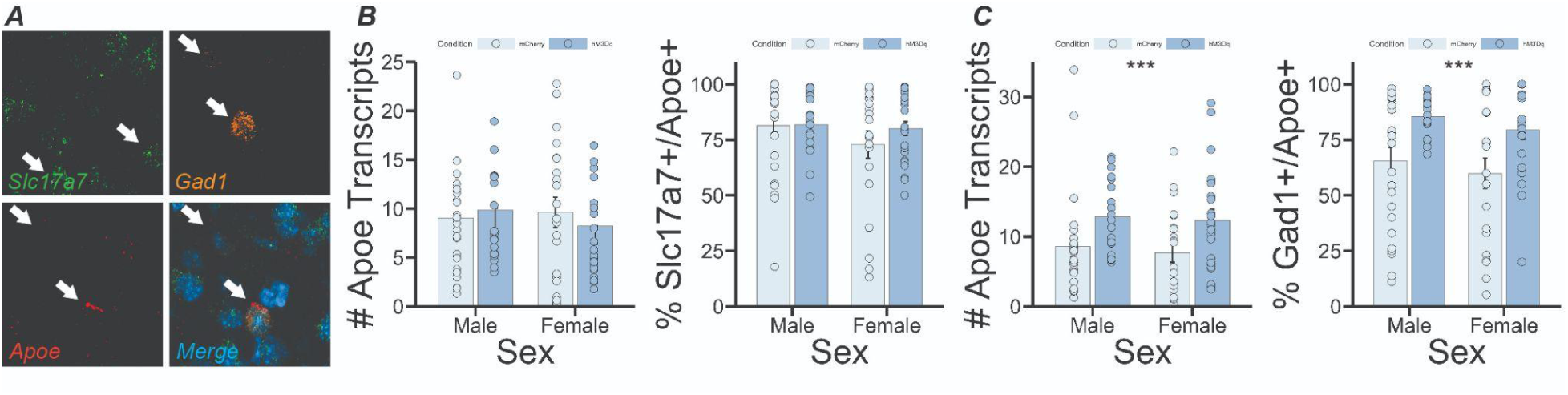
(A) Confocal image showing expression of Slc17a7, Gad1, and Apoe in FC tissue. (B) No significant differences in Apoe expression in glutamatergic cells were observed between conditions or sexes. (C) Apoe expression significantly increased in GABAergic neurons following LC depolarization (F(1,84) = 9.154, p = 0.00329, main effect of condition for # transcripts, F(1,84) = 12.898, p = 0.000553, main effect of condition for % GABAergic neurons).

Finally, we looked at *Apoe* expression in two different subtypes of inhibitory neuron - parvalbumin interneurons (*Pvalb*-expressing cells), and somatostatin interneurons (*Sst-expressing* cells; Fig. 5a). We found no evidence of increased *Apoe* expression in putative parvalbumin interneurons in the experimental group - however, we did find an increased number of *Apoe* transcripts in *Pvalb*-expressing cells in females compared to males (*F*(1,86) = 12.961, *p* = 0.000531, main effect of sex), as well as an increase in the percentage of *Pvalb*-expressing cells that co-expressed *Apoe* in females compared to males (*F*(1,86) = 5.536, *p* = 0.0209; Fig. 5b), suggesting that there are baseline differences in *Apoe* expression in cortical parvalbumin interneurons between sexes in mice. *Apoe* expression was, however, increased in *Sst*-expressing cells in the experimental group (*F*(1,86) = 11.088, *p* = 0.001281, main effect of group for number of *Apoe* transcripts). The increase in *Apoe* expression in putative somatostatin interneurons was partially sex-dependent - number of *Apoe* transcripts in *Sst*-expressing cells was also higher in females compared to males (*F*1,86) = 13.425, *p* = 0.000429, main effect of sex for number of *Apoe* transcripts), and analysis of the percentage of *Sst*-expressing cells that co-expressed *Apoe* revealed a sex x group interaction (*F*(1,86) = 7.476, *p* = 0.00759), with an increase in the female experimental group specifically.

**Fig. 5:**
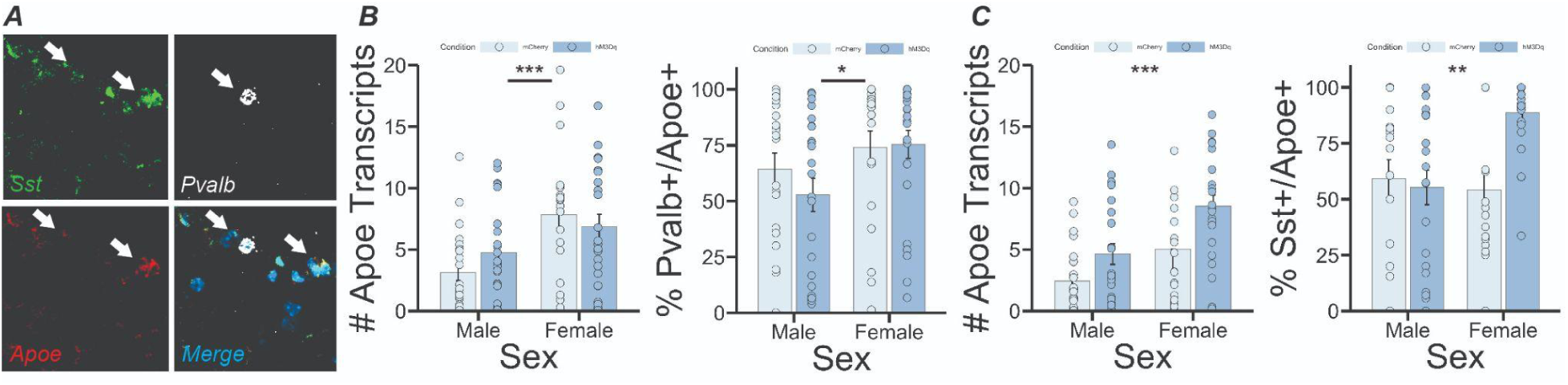
(A) Confocal image showing expression of Pvalb, Sst, and Apoe in FC tissue. (B) Apoe expression in parvalbumin interneurons was significantly higher in females (F(1,86) = 12.961, p = 0.000531, main effect of sex for # transcripts, F(1,86) = 5.536, p = 0.0209, main effect of sex for % parvalbumin interneurons). (C) Apoe expression significantly increased in somatostatin interneurons following LC depolarization (F(1,86) = 11.088, p = 0.001281, main effect of condition, F(1,86) = 13.425, p = 0.000429, main effect of sex for # transcripts, F(1,86) = 7.476, p = 0.00759, condition x sex interaction for % neurons).

## Discussion

We find that chemogenetic activation of a circuit that is involved in attention (Hallock et al., 2024) induces cortical expression of genes in mice that could be used for molecular identifiers of function in this circuit. Specifically, we find that one of these genes (*Apoe*) is selectively enriched in GABAergic neurons following circuit activation. These results provide insight into how distinct anatomical inputs might regulate the transcriptome of cortical regions that are involved in a variety of behaviors, such as the frontal cortex. The design employed in this study compliments research demonstrating behaviorally-induced (Cho et al., 2016; Mukherjee et al., 2018) and stimulation-induced (Nelson et al., 2023; Bach et al., 2024) changes in gene expression in rodents. The identification of genes that are upregulated upon activation of attentional circuits might serve as a starting point for the development of targeted treatments for attentional deficits in disorders such as ADHD, schizophrenia, and MDD.

The *APOE* gene encodes for a polymorphic protein that is implicated in processes such as neurogenesis, plasticity, and neuronal repair (Rusted et al., 2013). *APOE* has four allelic variants in humans, with the fourth allele (-e4) being heavily linked to the development of Alzheimer’s Disease (AD). Patients with AD exhibit many facets of attentional deficits; for example, selective attention, or the ability to attend to a particular stimulus in the environment, and executive control of attention, or coordinating goal-directed activities, are both impaired in AD (Parasuraman et al., 2002). *APOE* genotype, even in the absence of AD pathology, modulates a range of processes, including visuospatial attention and working memory (Espeseth et al., 2006). Interestingly, deficits in attention, specifically the shifting of visuospatial attention, have been found in *APOE* -e4 allele carriers who do not have dementia (Greenwood et al., 2000).

*APOE* allele type impacts facets of attention across the lifespan as well; younger individuals who are carriers of the -e4 allele perform better on measures of sustained and covert attention when compared to younger individuals who are not carriers of the -e4 allele (Rusted et al., 2013). Interestingly, advantages in performance noted early in life for -e4 carriers might turn into disadvantages later in life. Filippini et al. (2010) looked at blood-oxygen level dependent (BOLD) signals in both older and younger carriers, and non-carriers, of the -e4 allele. They found that older -e4 individuals exhibited significantly increased activation of several brain regions relative to younger -e4 carriers, an effect that was not observed between young and old -e4 carriers.

Filippini et al. (2010) proposed that the -e4 allele might impact age-related compensatory processes. It is possible that the age- and allele-related decline in task performance observed with relation to *APOE* is related to neurophysiological changes, such as an increase in amyloid beta deposition, which is a hallmark of AD (Filippini et al., 2010). The -e4 allele has the least amount of lipidation out of any of the *APOE* alleles (Flowers & Rebeck, 2020), which means that the proteins encoded by the *APOE* -e4 allele are more likely to aggregate and become toxic to neurons.

In order to fully understand the function of *APOE* as it relates to attention and disease state, it is necessary to know where *APOE* is being expressed within attentional circuitry. Attentional deficits are becoming more recognized as a hallmark of AD, as they appear before memory loss (Parasuraman et al., 2002). LC neurons also degenerate over the progression of AD, and have been identified as one of the first locations in the brain to accumulate tau protein, a pathological hallmark of AD (James et al,, 2020; Dahl et al., 2020). Ideally, if *APOE* is targeted as a potential treatment for attentional deficits, this treatment might serve to alleviate issues that occur early on in the progression of AD. In a healthy human brain, *APOE* is expressed primarily in astrocytes (Flowers & Rebeck, 2020). Astrocytes provide neurons with cholesterol, and *APOE* is the predominant carrier for cholesterol transport to these neurons (Li et al., 2022). Similar to humans, *Apoe* is primarily expressed in astrocytes in wild-type rodent brains (Raber et al., 1998). However, Xu et al. (2006) found that upon excitotoxic injury, hippocampal neurons in the mouse brain will express *Apoe*. Xu et al. (2006) also proposed that, depending on where *Apoe* is being expressed, it could serve different functions; for example, in damaged neurons, *Apoe* might be involved in mitochondrial dysfunction and neurofibrillary tangle formation, and in astrocytes, it may be involved in the formation of amyloid plaques. Other papers have demonstrated that *APOE* is expressed in human (Xu et al., 1999) and rodent (Boschert et al., 1999; Harris et al., 2004) neurons, suggesting that the APOE protein functions in both astrocytes and neurons to regulate circuit function. Elucidating where *Apoe* is being expressed in brain regions that are involved in attention may shed light on what *Apoe*’s role is in those circuits, and which metabolic products or molecules related to neuropsychiatric conditions *Apoe* is involved in producing. Our finding that activation of one such attentional circuit induces upregulation of *Apoe* in neurons, and not astrocytes, is a step toward this goal.

We found that there were no statistically significant differences in the average number of *Apoe* transcripts per putative excitatory (*Slc17a7*-expressing) neuron, as well as the proportion of *Slc17a7*-expressing neurons that co-expressed *Apoe* between DREADD-expressing and control mice. These results suggest that excitatory neurons are therefore not responsible for the upregulation of *Apoe* found upon activation of LC-FC projection neurons. We instead observed that a significantly greater average number of inhibitory neurons co-expressed *Apoe* in DREADD-expressing mice, and that the average number of *Apoe* transcripts in *Gad1*-expressing cells was significantly higher in DREADD-expressing mice compared to controls. We therefore conclude that inhibitory neurons are responsible for the upregulation of *Apoe* following depolarization of FC-projecting neurons in the LC. Inhibitory neurons are involved in sculpting local networks; they mostly lack long range projections, and instead influence the activity of local circuitry (Moore, 1993). The major neurotransmitter used by inhibitory neurons is gamma-aminobutyric acid (GABA). In the present study, we used *Gad1* as a general marker for inhibitory neurons. The *Gad1* gene encodes for glutamic acid decarboxylase (GAD), which is an enzyme that catalyzes the decarboxylation of glutamate in GABA synthesis (Bu et al., 1992).

Previous research suggests that there is increased susceptibility of GABAergic interneurons to *APOE* -e4 related pathology (Najm et al., 2019). There exists an interesting connection between *APOE* allele type and GABAergic inhibitory activity, in that *APOE* -e4 has been associated with hyperactivity of brain function, measured through BOLD signals, in young carriers (Filippini et al., 2010). It is possible that this reflects a difference in GABAergic inhibitory neuronal function, especially given the association between *APOE* -e4 and subclinical epileptiform activity that can occur in patients without dementia when experiencing stress (Palop & Mucke, 2009; Andrews-Zwilling et al., 2010). At a molecular level, *APOE* -e4 undergoes proteolytic cleavage in neurons, which generates neurotoxic fragments (Brecht et al., 2004). These fragments eventually lead to tau-phosphorylation, a key component, alongside amyloid beta plaque formation, of AD (Brecht et al., 2004; Najm et al., 2019). Li et al. (2009) found that levels of neurotoxic *Apoe* fragments and tau phosphorylation were elevated in the hippocampal neurons of mice that expressed *APOE* -e4. They also found decreased levels of GABAergic interneuron survival in these mice. This finding demonstrates that GABAergic interneurons, specifically in brain regions like the hippocampus, are susceptible to *APOE* -e4-derived neurotoxic fragments and tau pathology (Andrews-Zwilling et al., 2010). Both factors contribute to decreased survival of GABAergic interneurons, which also leads to learning and memory deficits characteristic of AD (Knoferle et al., 2014).

Because *Gad1* is a general marker for inhibitory neurons, it cannot delineate which subtype of interneuron, if any, is responsible for the upregulation of *Apoe* found in this experiment. To investigate this, we examined *Apoe* expression in two major subtypes of inhibitory neuron: parvalbumin interneurons and somatostatin interneurons. Cortical parvalbumin interneurons mainly provide peri-somatic inhibition to excitatory pyramidal neurons, while somatostatin interneurons mainly target pyramidal neuron dendrites (Hangya et al., 2014). We find that enrichment of *Apoe* is primarily observed in putative somatostatin interneurons following activation of FC-projecting LC neurons. Furthermore, this effect is much stronger in female mice, revealing that cell type-specific expression of *Apoe* is sex-dependent. Somatostatin interneurons in cortex are widely implicated in a range of behaviors that are affected in neuropsychiatric disorders, such as working memory (Kim et al., 2016; Abbas et al., 2018), attention (Urban-Ciecko et al., 2018), and learning (Adler et al., 2019). *Apoe* may therefore act as a molecular regulator of function in these neurons during attentional processing. Interestingly, previous research has demonstrated that the distribution of somatostatin interneurons in several brain regions is sexually dimorphic (Kim et al., 2017). Deficits in somatostatin gene production in the cingulate cortex of patients with MDD is also more pronounced in female patients (Tripp et al., 2011; Seney et al., 2015), indicating that sex differences in somatostatin interneuron function may contribute to differences in the onset and severity of neuropsychiatric disorders between biological males and females. Further research is needed to uncover which interneuron subtypes underlie increases in *Apoe* expression observed in male mice in our study.

Taken together, our findings provide an interesting link between inhibitory neurons and activity-induced *Apoe* expression in the frontal cortex. Although the research surrounding *Apoe* and GABAergic interneurons has traditionally focused on learning and memory deficits and allele-specific effects, it is possible that a connection exists between *Apoe*, GABAergic interneuron function, and attentional deficits. Loss of GABAergic interneurons in *APOE* -e4 carriers is associated with learning and memory deficits, but these occur after attentional deficits arise during the progression of AD. Therefore, it is possible that *APOE*-related GABAergic interneuron loss is occurring in brain regions that are involved in attention, such as the FC and the LC. Our results suggest that cell type-specific regulation of *Apoe* might be a therapeutic strategy for treating attention symptoms in neuropsychiatric and neurodegenerative disorders. Importantly, although the structure of the mouse *Apoe* gene and human *APOE* gene are different, a basic understanding of how and where *Apoe* is expressed in specific circuits is a foundational step toward illuminating how human alleles might impact cellular function to affect cognitive domains such as attention.

## Author Contributions

GEC designed research, performed research, and wrote the paper; LR performed research; SRE performed research; NJE analyzed data; AEJ designed research and analyzed data; KM designed research; HLH designed research, analyzed data, and wrote the paper

## Acknowledgments

The authors would like to acknowledge Aimee Ormond, Deveren Manley, and Amy Badillo for animal care assistance

